# Functional genomics screens reveal a role for TBC1D24 and SV2B in antibody-dependent enhancement of dengue virus infection

**DOI:** 10.1101/2024.04.26.591029

**Authors:** Laura Belmont, Maya Contreras, Catiana H. Cartwright-Acar, Caleb D. Marceau, Aditi Agrawal, Lisa M. Levoir, Jay Lubow, Leslie Goo

**Affiliations:** Vaccine and Infectious Disease Division, Fred Hutchinson Cancer Center, Seattle, Washington, USA; Molecular and Cellular Biology Graduate Program, University of Washington, Seattle, Washington, USA; Chan Zuckerberg Biohub, San Francisco, California, USA

## Abstract

Dengue virus (DENV) can hijack non-neutralizing IgG antibodies to facilitate its uptake into target cells expressing Fc gamma receptors (FcgR) - a process known as antibody-dependent enhancement (ADE) of infection. Beyond a requirement for FcgR, host dependency factors for this non-canonical infection route remain unknown. To identify cellular factors exclusively required for ADE, here, we performed CRISPR knockout screens in an *in vitro* system permissive to infection only in the presence of IgG antibodies. Validating our approach, a top hit was FcgRIIa, which facilitates binding and internalization of IgG-bound DENV but is not required for canonical infection. Additionally, we identified host factors with no previously described role in DENV infection, including TBC1D24 and SV2B, both of which have known functions in regulated secretion. Using genetic knockout and *trans*-complemented cells, we validated a functional requirement for these host factors in ADE assays performed with monoclonal antibodies and polyclonal sera in multiple cell lines and using all four DENV serotypes. We show that knockout of TBC1D24 or SV2B impaired binding of IgG-DENV complexes to cells without affecting FcgRIIa expression levels. Thus, we identify cellular factors beyond FcgR that are required for ADE of DENV infection. Our findings represent a first step towards advancing fundamental knowledge behind the biology of ADE that can ultimately be exploited to inform vaccination and therapeutic approaches.

## 1. Introduction

The complicated antibody response to the four circulating serotypes of dengue virus (DENV1-4) represents a major barrier to the development of safe and effective vaccines and therapeutics. Specifically, primary infection with one DENV serotype does not confer durable immunity against infection by the other three serotypes. Instead, the biggest risk factor for dengue disease is secondary infection with another DENV serotype in the presence of pre-existing DENV-specific IgG antibodies from a prior exposure [^1–5^]. The prevailing theory behind this phenomenon of antibody-dependent enhancement (ADE) is that non-neutralizing IgG antibodies facilitate virus uptake into target cells through Fc-Fc gamma receptor (FcgR) interactions[^6^].

*In vitro* studies have established that antibody-mediated neutralization and enhancement of infection depends on IgG concentration [^7^]. This is evident in the infectivity curve observed in FcgR+ cell lines (such as K562 and U937) widely used to study ADE as they are permissive to infection only in the presence of DENV-reactive IgG [^8,9^]. In these cells, peak enhancement of infection occurs at intermediate antibody levels; higher antibody concentrations neutralize infection, while lower concentrations do not enhance infection. Importantly, the risk of severe dengue disease risk in humans is also highest within a narrow range of intermediate titers of pre-existing antibodies to DENV[^1,2,5^].

Canonical DENV infection in the absence of antibodies is predominantly initiated via classical clathrin-dependent endocytosis following direct virion interaction with cellular attachment factors [^10–13^]. In contrast, efficient uptake of IgG-opsonized DENV is dependent on intact signaling of FcgRs such as FcgRIIa [^8,14^], an activating FcgR expressed on relevant DENV target cells *in vivo* [^9,15–20^]. Live-cell imaging and single-particle tracking studies found that actin-mediated plasma membrane protrusions facilitated the uptake of IgG-opsonized but not ‘naked’ DENV particles [^21^], suggesting that unique entry factors are important for ADE. Additionally, IgG-dependent uptake of DENV particles can increase not only the number of infected cells, but also the viral output per infected cell [^6^], suggesting that DENV-host interactions downstream of viral entry also contribute to distinct infection outcomes observed under ADE and non-ADE conditions. This is unsurprising given that Fc-FcgR signaling regulates many cellular processes, including endocytosis, cell proliferation and maturation, and innate immunity [^22^].

Beyond dependence on antibody concentration and the role of FcgR in initiating uptake of IgG-bound virions, the functional requirements for DENV infection via ADE are still unknown. This knowledge gap persists largely because *in vivo* models fail to fully capture DENV immunity and pathogenesis [^23–25^]. Our limited understanding of potential ADE mechanisms comes from *in vitro* studies that do not establish cause and effect. Most previous studies selectively investigated the expression of specific cytokines in response to IgG-mediated entry of DENV [^9,26–33^]. Conversely, unbiased transcriptomic profiling of whole blood or peripheral blood mononuclear cells have identified differentially expressed genes in patients with mild versus severe dengue disease [^27,34,35^], but these studies cannot establish whether the observed profiles are specific to antibody-mediated infection. Although a recent study showed that DENV infection via ADE uniquely altered the expression of multiple host genes [^36^], their direct functional contribution to ADE was not explored. Thus, existing studies have only indirectly suggested potential ADE mechanisms.

A comprehensive analysis of which host factors are functionally required for ADE is lacking. Genome-wide CRISPR knockout screens have enabled high-throughput, unbiased, and reproducible discovery of viral host dependency factors [^37^]. Such screens of direct (non-ADE) DENV infection in the absence of antibodies performed independently by multiple researchers using multiple cell lines and viral strains have identified highly concordant host dependency factors [^38–42^]. However, as these prior screens were performed in the context of non-IgG-mediated DENV infection, they could not identify host factors uniquely required for ADE.

Here, to advance mechanistic understanding of ADE beyond existing descriptive studies, we performed a CRISPR/Cas9-based genome-wide and follow-up targeted knockout screens in K562 cells, which are permissive to efficient infection only via ADE [^14^]. This approach was designed to reveal host factors exclusively required for ADE mechanisms. Validating this approach, our screens identify candidate ADE-specific host factors with no previously defined role in DENV infection, including TBC1D24 and SV2B, both of which are essential in trafficking specialized recycling endosomes during regulated secretion [^43,44^]. We validated a functional role for TBC1D24 and SV2B in promoting ADE of all four DENV serotypes, with monoclonal antibodies and polyclonal sera, and in multiple cell lines. Further, we show that knockout of TBC1D24 or SV2B reduced the efficiency of binding of IgG-DENV complexes to cells despite maintaining similar levels of FcgRIIa expression to unedited cells.

Thus, we identify for the first time host factors beyond FcgR that are required for efficient ADE of DENV infection. Our screen represents a novel discovery tool for cellular factors and processes uniquely subverted by DENV during ADE that can be exploited to significantly advance our understanding of ADE mechanisms.

## 2. Materials and Methods

### 2.1. Cell lines

K562 cells (Cat# CCL-243, ATCC), U937 cells (provided by Taia Wang, Stanford University), and Raji cells stably expressing DCSIGNR (Raji-DCSIGNR) (provided by Ted Pierson, NIAID, NIH, Bethesda, MD) were maintained in RPMI 1640 supplemented with GlutaMAX (Cat# 72400-047; ThermoFisher Scientific), 7% fetal bovine serum (FBS) (Cat# 26140079, lot 2358194RP, ThermoFisher Scientific) and 100 U/mL penicillin-streptomycin (Cat# 15140–122; ThermoFisher Scientific). HEK-293T/17 cells (Cat# CRL-11268, ATCC) were maintained in DMEM (Cat #11965118; ThermoFisher Scientific) supplemented with 7% FBS and 100 U/mL penicillin-streptomycin. C6/36 cells (Cat # CRL-1660, ATCC) were maintained in EMEM (Cat # 30-2003, ATCC) supplemented with 10% FBS. TZM-bl cells (provided by Michael Emerman, Fred Hutchinson Cancer Center, Seattle, WA) were maintained in DMEM (Cat #11965118; ThermoFisher Scientific) supplemented with 7% FBS and 100 U/mL penicillin-streptomycin.

K562-DCSIGN cells were generated by lentiviral transduction. A plasmid expressing DCSIGN (Genbank Accession NM_021155.4) fused to BFP in a lentiviral vector was synthesized (VB221014-1121vtg, VectorBuilder) and used to transduce K562 cells as described below in “Lentiviral production and transduction.” Transduced cells were stained using anti-DCSIGN antibody (Cat #330105, Biolegend) and cell populations highly expressing both DCSIGN and BFP were bulk sorted (Sony MA900).

C6/36 cells were maintained at 30°C in 5% CO2; all other cell lines were maintained at 37°C in 5% CO2.

### 2.2. Viruses

DENV1 UIS 998 (isolated in 2007, Cat# NR-49713), DENV2 US/BID-V594/2006 (isolated in 2006, Cat# NR-43280), DENV3/US/BID-V1043/2006 (isolated in 2006, Cat# NR-43282), and DENV4 strain UIS497 (isolated in 2004, Cat# NR-49724) were obtained from BEI Resources (Manassas, VA) and propagated on C6/36 cells. Virus-containing supernatant from days 3 to 8 post-infection was pooled, centrifuged at 500 x *g* for 5 min, filtered through a 0.45 μm Steriflip filter (Cat# SE1M003M00, Millipore-Sigma), and stored at -80°C until use. DENV2 S16803 reporter virus particles (DENV2-GFP) (Cat # RVP-201) [^45^] were purchased from Integral Molecular, Inc (Philadelphia, PA). Infectious titers of viral stocks were determined by infecting Raji-DCSIGNR cells with 2-fold serial dilutions of virus. Cells infected with fully infectious virus were fixed and permeabilized using BD cytofix/cytoperm (Cat #554717, BD Biosciences) according to the manufacturer’s instructions, stained with APC-conjugated E-protein-specific antibody, 4G2 for 30 minutes at 4°C and washed in cytoperm/wash buffer twice before quantification of APC+ cells by flow cytometry (Intellicyt iQue Screener PLUS, Sartorius AG). Antibody 4G2 was isolated and purified by the Fred Hutchinson Cancer Center Antibody Technology core following expansion of the hybridoma D1-4G2-4-15 (Cat# HB-112, ATCC) and purification of IgG from culture supernatant. Purified 4G2 was conjugated to APC via the Lighting-Link APC-conjugation kit (Cat# ab201807, Abcam) according to manufacturer’s instructions. Cells infected with DENV2-GFP were fixed with 2% paraformaldehyde and GFP+ cells quantified by flow cytometry (Intellicyt iQue Screener PLUS, Sartorius AG).

### 2.3. Lentiviral production and transduction

Lentivirus was produced in HEK-293T/17 cells by co-transfection of lentiviral plasmids with psPAX2 (Cat# 12259, Addgene) and pMD2.G (Cat# 12259, Addgene) at a mass ratio of 2:1.33:1, respectively, using the Lipofectamine™ 3000 Transfection Reagent (Cat# L3000001, ThermoFisher Scientific). Supernatants were collected 48 hours post-transfection, passed through a 0.22 μM filter, and either stored at -80°C or immediately used to transduce cells.

Target cells were seeded in 6-well plates at a density of 2e5 cells per well in 2 mL RPMI 1640 with 7% FBS and 1% penicillin-streptomycin the day prior to transduction. On the day of transduction, cells were pelleted and resuspended in 250 μL lentivirus and 8 μg/mL DEAE-Dextran (Cat# D9885, Sigma) in a total volume of 1.7 mL, followed by spinoculation at 1000 x *g* for 2h at 30°C. Medium on spinoculated cells was aspirated and replaced with 2 mL fresh RPMI formulated as above. After incubation at 37°C for 24h, cell culture medium was replaced again. After at least 3 days since spinoculation, transduced cells were either bulk sorted by FACS (Sony MA900) or subjected to drug selection, depending on lentivirus vector marker.

### 2.4. Genome-wide and targeted CRISPR screens

K562-Cas9-Blast cells (provided by Andreas Puschnik, Chan Zuckerberg Biohub, San Francisco) were generated by transduction with lentiCas9-Blast (Cat# 52962, Addgene) and selection with blasticidin as described previously [^38^]. Approximately 200 million K562-Cas9-Blast cells were transduced with each GeCKOv2 library A or B [^46^] (Addgene #1000000048 or #1000000049, respectively) at a MOI of 0.3 in the presence of 10 μg/mL protamine sulphate. Three days post-transduction, cells were selected in 1 μg/mL puromycin. Sixty four million mutagenized cells for each library (A and B) were infected with DENV2-GFP at an MOI of 24 in the presence of 1.25 μg/mL anti-DENV2 mouse antibody DV2-70 [^47^] (provided by Michael Diamond, Washington University, St. Louis, MO) by spinoculation at 930 x g for 2 hours at 30°C, and then incubated at at 37°C for two days. Next, to isolate ADE-resistant cell populations, GFP-negative cells were bulk sorted (Sony SH800) and allowed to recover and multiply for five days at 37°C prior to re-infection by ADE under the above conditions. After three total rounds of ADE and bulk sorting, genomic DNA was isolated using the QIAamp DNA Blood Mini Kit (Cat #51185, Qiagen), and sgRNA sequences were amplified and prepared for next-generation sequencing via the NextSeq platform (Illumina). The enrichment of each sgRNA in the selected cells relative to the unselected libraries cultured and harvested in parallel was calculated using MAGeCK [^48^].

The custom targeted library against the top 500 highest-ranking candidates from the genome-wide screen was designed using CHOPCHOP v3 and Guides [^49,50^]. We used six guides per gene, plus 200 non-targeting control (NTC) guides (**Table S2**). Pooled gRNAs were synthesized (Twist Biosciences, South San Francisco, CA) and cloned into lentiCRISPRv2 (Cat# 52961, Addgene), packaged into lentivirus as described above, and titered on TZM-bl cells, as previously described [^51^]. Approximately 6.4 million K562 cells were transduced with the targeted lentivirus library at an MOI of 0.5 (500-fold coverage) in the presence of 8 μg/mL of DEAE-Dextran by spinoculation at 930 x g for 2 hours at 30°C. The following day, cells were selected for two weeks using 1 μg/mL of puromycin. Mutagenized cells (3.2 million) were subjected to three rounds of infection via ADE and sorting, and subsequently prepared for next-generation sequencing (Illumina MiSeq) and analysis as described above for the genome-wide screen. Targeted screens were performed in biological triplicate on three independently mutagenized cell library populations.

### 2.5. Generation of clonal KO cell lines

The following individual KO cell lines were generated by nucleofection of Cas9-sgRNA ribonucleoprotein complexes (RNPs): K562 (TBC1D24 KO), U937 (TBC1D24 KO and SV2B KO), and K562-DCSIGN (NTC, TBC1D24 KO, FcgRIIa KO). RNPs were assembled by combining 6 μL of 30 pmol/μL of a pre-made mixture of three equimolar sgRNAs (Synthego Gene KO Kit v2), 1 μL of 20 uM Cas9-NLS protein (Synthego), and 18 μL of SF Cell Line Nucleofector solution (Lonza V4XC-2032). Complexes were mixed and incubated for 10 min at room temperature prior to addition to 5 μL of cells (1e5 for U937, 2e5 for K562). This mixture was transferred to a 16-well nucleocuvette for nucleofection with an Amaxa 4D Nucleofector (Lonza) according to manufacturer’s protocol using pulse code FF-120 for K562 cells or EP-100 for U937 cells. Following nucleofection, cells were incubated for 10 minutes at room temperature, and then transferred into a 24-well plate in a total volume of 500 μl for recovery. After 24 hours, the medium on the cells was replaced. Seventy two hours post-nucleofection, single clones were isolated via limiting dilution.

SV2B K562 KO, FcgRIIa K562 KO, and SV2B K562-DCSIGN KO cell lines were generated via nucleofection with top-ranking sgRNAs (**Table S4**) cloned into px458 (Cat# 48138, Addgene). One million cells resuspended in 100 μL of SF Cell Line Nucleofector solution (Lonza V4XC-2012) were electroporated with 2 μg plasmid using pulse code FF-120 (Lonza Amaxa 4D Nucleofector). Cells were incubated for 10 minutes at room temperatue, resuspended in 500 μL of RPMI with 7% FBS and 1% penicillin-streptomycin, and then moved into a 6-well plate for recovery at 37*°C.* After 48 hours, GFP-positive single cells were sorted into 96-well plates (Sony SH800).

For genotyping, genomic DNA was isolated (QuickExtract, Cat #QE0905T, Lucigen) and the gRNA-targeted site was amplified by PCR (primers listed in **Table S4**) and Sanger sequenced. Reads were aligned to reference sequences obtained from parental unedited (WT) cells and analyzed for the presence of indel mutations (Geneious Prime 2020.1.2). Mixed traces from heterozygous clones were deconvolved using ICE analysis [^52^].

FcgRIIa KO U937 cells have been previously described [^35^] and were provided by Taia Wang (Stanford University).

### 2.6. Generation of trans-complemented cell lines

Lentiviral transduction as described above was used to generate *trans-*complemented KO lines. The following cDNA constructs were obtained: SV2B (Cat #OHu12014D, GenScript) and TBC1D24 (Cat #OHu21983, GenScript). The cDNA of interest was amplified by PCR using primers (**Table S4)** with overhangs that allow directional cloning into EcoRV-linearized pHIVdTomato (#21374, Addgene) using the 2X Gibson Assembly Kit (Cat # E2611S, New England Biolabs). Assembled constructs were confirmed by whole plasmid sequencing (Plasmidsaurus, Eugene, OR). KO cells were transduced with lentiviral preparations to deliver the gene of interest and then bulk sorted (Sony MA900) based on high dTomato expression.

### 2.7. Validation ADE assays

Dose-dependent ADE assays were performed with fully infectious versions of DENV1-4 or single-round infectious DENV2-GFP reporter virus particles. Viral stocks were diluted to 5-10% final infectivity (determined on Raji-DCSIGNR cells as described above) and incubated with 5-fold serial dilutions of monoclonal antibody or polyclonal sera for 1 hr at room temperature before addition of 2e5 (in a 384-well plate) or 3.33e5 (in a 96-well plate) K562 or U937 cells, respectively. DV2-70 mouse monoclonal antibody was provided by Michael Diamond (Washington University, St. Louis, MO) [^47^] and human monoclonal J9 IgG was recombinantly produced as previously described [^53^]. Human convalescent serum samples from three independent DENV-immune donors were obtained from BEI Resources (Cat #NR-50229, NR-50232, NR-50231). Following incubation at 37°C for 2 days, cells were processed according to the protocol described above (section “Viruses”) and infection was quantified by flow cytometry (Intellicyt iQue Screener PLUS, Sartorius AG).

### 2.8. Direct infection of K562-DCSIGN cells

Two hundred thousand cells in 20 μL complete RPMI were infected with an equal volume of DENV2-GFP at a MOI of 24 and added in duplicate to a 384-well plate by spinoculation for 2 hours at 33°C and 1000 x*g*. Cells were resuspended and incubated at 37°C for 2 days prior to quantification of infected cells by flow cytometry as described above.

### 2.9. RT-qPCR infection assays

DENV2-GFP stocks at an MOI of 24 were incubated with 80 ng/mL J9 antibody for 1 hour at 4°C before addition to 3.33e5 K562 cells in duplicate wells of a 96-well plate on ice; all components were at an equal volume of 33 μL. Virus/antibody complexes were allowed to bind to cells for 1 hr at 4°C, followed by wash steps with 1X PBS to remove unbound virus/antibody complexes. Next, cells were either immediately (0 h time point) harvested for quantitative PCR or incubated at 37°C for 15 min to trigger internalization. Following three wash steps in 1X PBS, cells were treated with 400 ng/μL proteinase K at 37°C for 45 minutes to remove non-internalized complexes. Following three wash steps with 1X PBS, cells were further incubated at 37°C for 2h, 6h, or 24h prior to lysis for quantitative PCR (QuantStudio^TM^ 5 Real-Time PCR System, 96-well, Applied Biosystems^TM^) using the Power SYBR™ Green Cells-to-CT™ Kit (Cat #4402954, Invitrogen) per manufacturer instructions. Data were analyzed using ABI QuantStudio 5 (Applied Biosystems). All viral RNA levels were normalized to 18S levels, and subsequently to control WT cells at 0 hours post-infection. Universal DENV primer and 18S primer sequences [^54^] can be found in **Table S4**.

### 2.10. Replicon assays

One million K562 cells were electroporated with 3 μg of DENV2-luciferase replicon [^38^] (provided by Jan Carette, Stanford University) in 100 μL SF Cell Line Nucleofector solution (Lonza V4XC-2012) using pulse code FF-120 (Amaxa 4D Nucleofector, Lonza). Cells were incubated for 10 minutes at room temperature following nucleofection, resuspended in 500 μL of RPMI with 7% FBS, then distributed into a 48-well plate (250 μL per well) and incubated at *37*°C. At each time point, cells were lysed using the Renilla-Glo® Luciferase Assay System (Cat #E2710, Promega) according to manufacturer suggestions, and frozen at -20°C. Samples from each timepoint were then concurrently processed using the Renilla-Glo® Luciferase Assay System according to manufacturer instructions, and analyzed on a plate reader (Infinite M1000 Pro, Tecan).

### 2.11. Determining FcgRIIa receptor expression

To assess surface FcgRIIa expression, 2e5 cells per cell type and stain were washed in FACS wash (FW, 2% FBS in 1x PBS) and resuspended in 50 μL anti-CD32-FITC (Cat # 60012.FI, StemCell) or isotype control (Cat # 11-4732-81, ThermoFisher Scientific), and incubated at 4°C for 20 min. FW was then added and a wash with FW was performed. Cells were then fixed in 2% PFA for 20 minutes, spun down, and resuspended in PBS at 4°C until acquisition. To assess total FcgRIIa expression, 2e5 per cell type and stain cells were fixed using Cytofix (Cat #554655, BD) for 20 minutes at 4°C cells before addition of perm/wash buffer (Cat #554723, BD) followed by two washes in perm/wash. Cells were then resuspended in either 50 μL anti-CD32-FITC or isotype control, and incubated at 4°C for 20 min. Cells were then washed with perm/wash prior to resuspension in PBS at 4°C until acquisition. Samples were analyzed via flow cytometry (Sony ID7000), and data were analyzed using FlowJo 10.9.0.

### 2.12. Statistical analysis

Area under the curve analysis, and paired and unpaired t-tests adjusted via Benjamini-Hochberg method were performed using GraphPad Prism 10.

## 3. Results

### 3.1. Genome-wide and targeted CRISPR screens identify host factors uniquely required for ADE

Our screening strategy is outlined in **Figure 1A**. To comprehensively identify candidate ADE-specific host dependency factors, we first generated a genome-wide knockout (KO) library of K562 cells. As mentioned, these cells are widely used to study ADE because they express FcgRIIa and are only permissive to DENV infection in the presence of IgG antibodies [^14^]. We infected the KO library with single-round infectious reporter virus particles of DENV2 strain S16803 (DENV2-GFP) [^45^] pre-complexed with DV2-70, a mouse DENV2-specific IgG antibody [^47^] under optimized conditions that achieved stringent selection pressure (>95% infection). To maximize signal-to-noise ratio, we performed three rounds of infection and used fluorescence-activated cell sorting (FACS) to isolate cells resistant to ADE and thus likely had host dependency factors knocked out. Specifically, after each round of infection via ADE, live, GFP-negative cells were sorted, allowed to recover and multiply, and then re-infected by ADE to ensure that lack of infection was due to gene KO instead of stochastic effects. We deep sequenced genomic DNA from the virus-selected cell population and used MAGeCK [^48^] to compare sgRNA enrichment relative to the uninfected KO library cultured and harvested in parallel.

**Figure 1.**
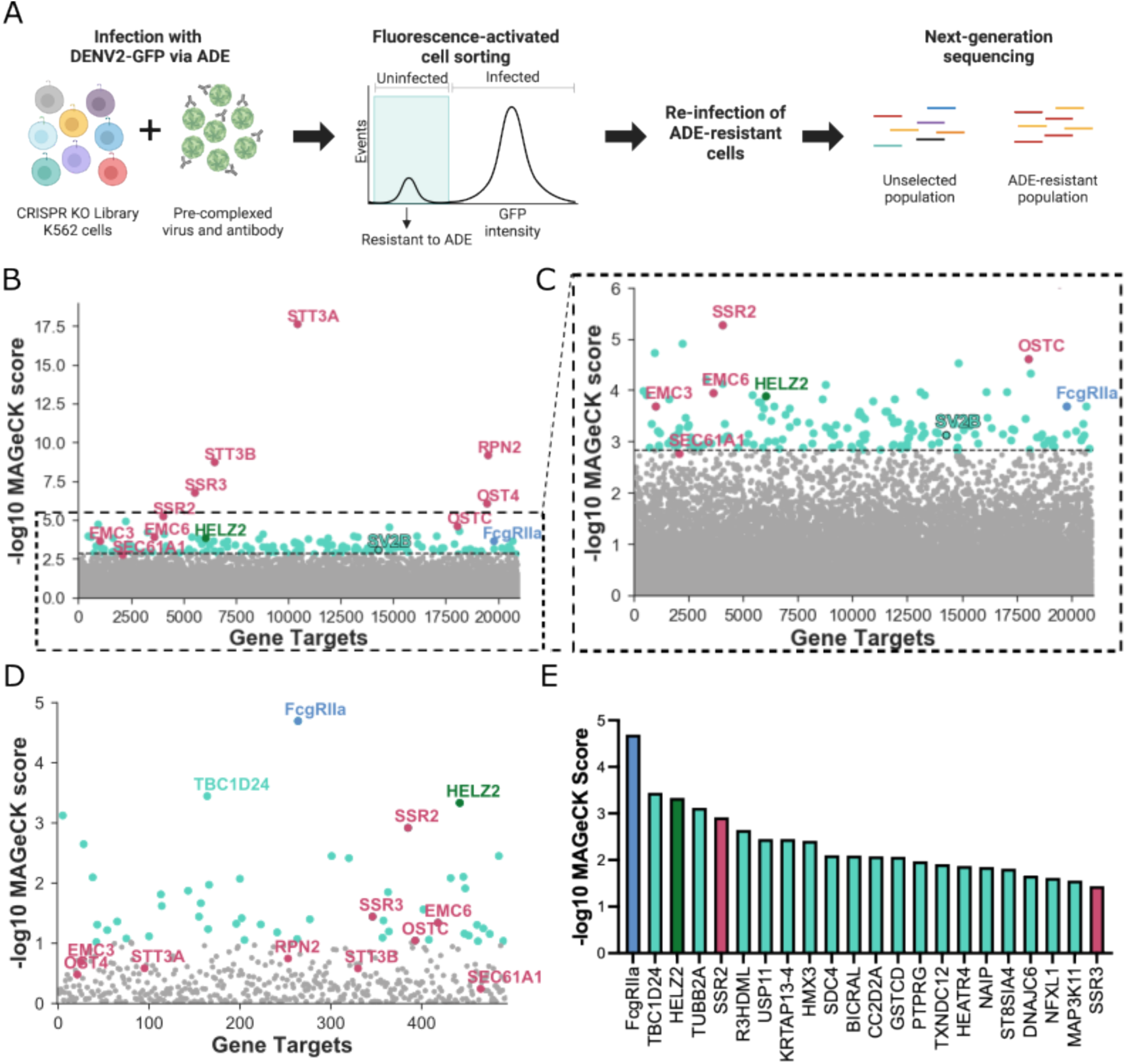
Genome-wide and targeted CRISPR knockout screens in human cells identify candidate host factors required for ADE of DENV infection**. (A)** Schematic for CRISPR-based genome-wide and targeted knockout screens. K562 cell mutant libraries were generated using either the GeCKOv2 sgRNA library or a compressed library targeting 500 genes for genome-wide or targeted screens, respectively. In both screens, cell libraries were infected via ADE with single-round infectious DENV2 expressing a GFP reporter (DENV2-GFP) pre-complexed with anti-DENV2 monoclonal IgG antibody. ADE-resistant cells were isolated via fluorescence-activated cell sorting, and re-infected with DENV2-GFP via ADE following recovery and expansion. After three iterative rounds of infection and sorting, we used deep sequencing and MAGeCK analysis to quantify the enrichment of sgRNAs in the infected cell population relative to the corresponding control uninfected population. **(B)** Gene enrichment in genome-wide CRISPR screen. Y-axis displays MAGeCK score in the infected cell population; x-axis displays gene targets arranged randomly. Light blue: FcgRIIa, a known host factor required for ADE. Maroon: pro-viral host factors identified in previous genome-wide screens in the context of direct DENV infection [^38–42^]. Green: HELZ2, a previously identified host factor that restricts direct infection [^55,56^]. Teal: candidate novel ADE-specific host factors, including SV2B. The dotted line intersects SEC61A1, the lowest-ranking of validated host dependency factors for direct DENV infection identified in previous screens [^70–72^]^;^ all genes below this point are shown in gray. **(C)** A zoomed-in view of graph shown in (**B)**, highlighting the enrichment of candidate novel ADE-specific host factors clustered around FcgRIIa. Color scheme is similar to (**B**). **(D)** Gene enrichment in targeted CRISPR screen. Color scheme is similar to (**B**), except that genes depicted in gray represent the bottom 90% ranking genes. **(E)** MAGeCK scores of top 23 genes enriched in the targeted screen. Color scheme is similar to (**B**). Panel (**A**) was created with Biorender.com.

As shown in **Figure 1B**, many highly-enriched hits include known host dependency factors identified in previous genome-wide screens in the context of direct (non-ADE) DENV infection [^38–42^]. In fact, the gene with the highest MAGeCK score by far was STT3A, which was identified in previous direct infection CRISPR screens and is required for efficient replication of DENV and other mosquito-borne flaviviruses [^38–42^]. This finding was unsurprising as we expect some shared features of direct infection and ADE. Interestingly, HELZ2, a previously described host factor that *restricts* direct flavivirus infection, was also enriched [^55,56^]. Validating our approach, FcgRIIa, a known ADE-specific host factor and the only activating FcgR expressed on K562 cells [^14^], was among the top hits. We also identified candidate ADE-specific genes like SV2B with no previously described role in DENV infection. In **Figure 1C**, we highlight that these novel genes were similarly enriched as FcgRIIa and interspersed among known direct infection host dependency factors identified in previous screens [^38–42^]. The full list of hits from the genome-wide screen is available in **Table S1**.

Although the enrichment of FcgRIIa indicated that our genome-wide screen was functioning as intended, many top hits were those identified in previous CRISPR screens in the context of direct infection [^38–42^] (**Figure 1B**). Therefore, to further enrich the most critical ADE-specific cellular factors, we performed a follow-up targeted screen using a custom sgRNA sub-library against the 500 highest-ranking genes from our genome-wide screen (**Tables S2-3**). To exclude potential off-target effects, we designed new sgRNA sequences distinct from those used in the initial genome-wide screen. As with the initial screen, K562 KO sub-libraries were infected with DENV2-GFP pre-complexed with DV2-70 IgG, followed by FACS of cells rendered resistant to ADE in three successive rounds. Unlike in the original genome-wide screen, the top hit in this targeted screen was our positive control, FcgRIIa (**Figure 1D**). Additionally, genes with no previously defined role in DENV infection dominated the top hits. These results demonstrate increased stringency in the targeted screen for identifying candidate ADE-specific factors over the genome-wide screen (compare **Figure 1D** to **Figures 1B-C**).

After FcgRIIa, the second top-ranking hit in the targeted screen was TBC1D24, which contains a Tre2/Bub2/Cdc16 (TBC) domain common to Rab-GTPase-activating proteins [^57^], and a TLDc domain with a putative function in oxidative stress resistance [^58^]. TBC1D24 is involved in recycling clathrin-independent endocytosis cargo proteins and in synaptic endocytic vesicle trafficking [^44,59^]. Interestingly, SV2B, a gene that scored highly in our genome-wide, but not targeted, screen (**Figure 1B-D**), is known to have related functions [^43,60–64^]. Specifically, SV2B is one of three paralogs of the SV2 family of integral membrane glycoproteins that regulate synaptic vesicle function via its role in trafficking synaptotagmin, a calcium sensor protein for exocytosis [^64,65^]. To our knowledge, neither TBC1D24 nor SV2B has a previously described role in virus infection. In fact, the top 23 hits of the targeted screen were dominated by novel candidate ADE-specific factors (**Figure 1E**). Like TBC1D24 and SV2B, some but not all high-ranking hits have known functions in the nervous system. These include TUBB2A, a microtubule component known to interact with KIF1a, which is required for synaptic vesicle transport [^66,67^]; DNAJC6, a heat-shock protein involved in neuronal clathrin-mediated endocytosis [^68^]; and HMX3, a transcription factor involved in neuronal cell specification [^69^].

### 3.2. Functional validation of TBC1D24 and SV2B as host dependency factors for ADE

We focused on validating the functional role of TBC1D24, the top-scoring gene in our targeted screen after FcgRIIa. We first generated TBC1D24 and FcgRIIa K562 KO clones and confirmed disruption of gene targets by PCR amplification and Sanger sequencing. FcgRIIa KO clone contained a frameshifting deletion (**Figure S1**) while TBC1D24 KO clone contained a large deletion at the beginning of exon 2, which is the first coding exon and encodes the TBC domain [^73,74^] (**Figure S2**). Next, we performed ADE dose-dependent assays using DENV2-GFP pre-complexed with mouse DV2-70 IgG, the same mouse antibody used in our screens. As expected, FcgRIIa KO largely abolished ADE (85% average reduction in area under the curve (AUC) compared to WT, **Figure S4**). Remarkably, TBC1D24 KO reduced ADE efficiency to similar levels as FcgRIIa KO (83% average reduction in AUC, **Figure S4A**).

Given the marked reduction of ADE efficiency due to TBC1D24 KO, we next investigated the role of SV2B, a host factor that shares a similar function as TBC1D24 in synaptic vesicle trafficking [^43,60–64^] and that was enriched in our genome-wide screen (**Figures 1B-C**). We generated a SV2B KO clone and confirmed a frameshifting deletion (**Figure S3)**. Dose-response ADE assays performed with this K562 SV2B-KO clone revealed a 70% reduction in AUC compared to WT (**Figure S4B**), thus demonstrating a functional role for SV2B in ADE.

To rule out a mouse antibody-specific artifact, we next performed the above ADE assays using a broadly reactive human monoclonal anti-DENV IgG antibody, J9 [^75^]. In these assays, TBC1D24 KO (**Figure 2A**) and SV2B KO (**Figure 2B**) each resulted in a ∼50% reduction in AUC compared to WT cells. Notably, ADE efficiency was rescued in KO cells *trans*-complemented with the gene of interest, but not with empty vector (**Figures 2A-B**), demonstrating that reduction in ADE efficiency was specifically due to KO of TBC1D24 or SV2B. Together, these experiments establish a functional role for TBC1D24 and SV2B in ADE.

**Figure 2.**
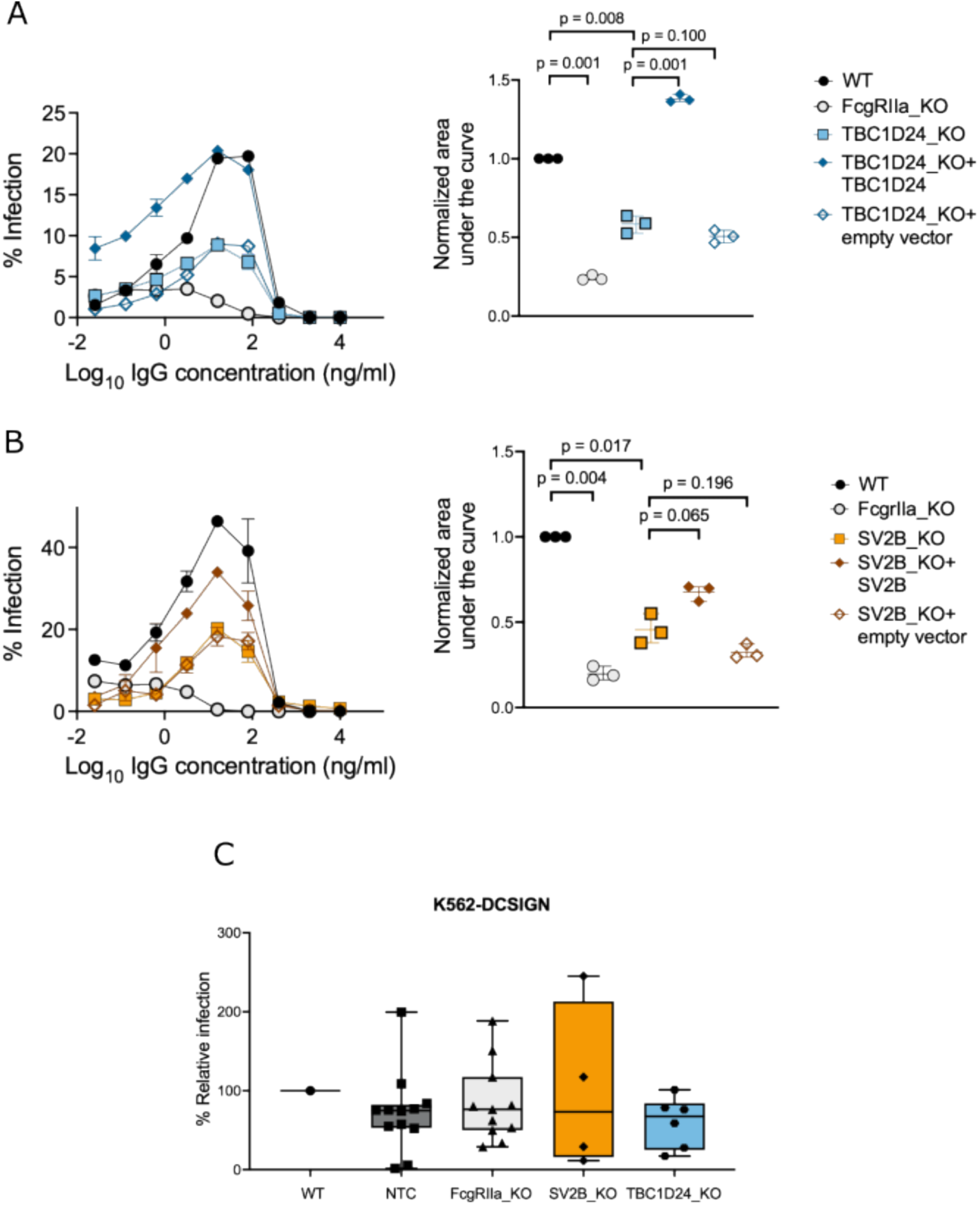
TBC1D24 and SV2B are required for efficient ADE but not direct infection. (**A-B**) (Left) Representative dose-response ADE curves for K562 (**A**) TBC1D24 KO clone (**B**) SV2B KO clone, and genetically *trans*-complemented K562 KO cells infected with DENV2-GFP in the presence of serially diluted human anti-DENV IgG monoclonal antibody J9 [^75^]. In each experiment, K562 WT cell pool and a FcgRIIa KO clone was included as a control. Data points and error bars represent the mean and range of infection in duplicate wells, respectively. Graphs shown are representative of at least three independent experiments. (Right) Quantification of area under the curve normalized to unmutagenized (WT) K562 cells from three independent dose-response ADE experiments, each represented as a data point. Horizontal lines and error bars indicate mean and standard deviation, respectively. P-values shown are from multiple paired student’s t-tests adjusted using the Benjamini-Hochberg method. (**C**) Efficiency of DENV2-GFP infection of the indicated K562-DCSIGN cells in the absence of IgG antibodies. Shown are percentage infections for individual KO clones (data points) normalized to unmutagenized (WT) K562-DCSIGN cell pool, median (horizontal line within box), 25th to 75th percentile (box), and minimum and maximum (whiskers). Data are representative of two independent experiments, each performed in duplicate wells. Comparisons of direct infection efficiency of each KO cell line to WT were not statistically significant (p > 0.05) as determined by multiple unpaired student’s t-tests adjusted using the Benjamini-Hochberg method.

Next, to confirm that the roles of TBC1D24 and SV2B are unique to IgG-mediated infection, we generated corresponding clonal KO lines in K562 cells engineered to express DCSIGN (K562-DCSIGN), a cellular attachment factor that permits direct DENV infection in the absence of IgG antibodies [^76^]. Following confirmation of target allele disruption (**Figures S5-S7**), we infected K562-DCSIGN KO clones with DENV2-GFP in the absence of antibodies. Due to substantial inter-clonal heterogeneity [^77^] even among non-targeting controls (NTC) (**Figure 2D**), we analyzed at least four KO clones per gene to mitigate clonal artifacts. FcgRIIa-KO, TBC1D24-KO, and SV2B-KO K562-DCSIGN clones reduced the efficiency of direct infection to relatively similar levels compared to the unedited (WT) K562-DCSIGN cell pool, (median reduction of 33%, 27%, 24%, respectively, **Figure 2D**). As FcgRIIa is required for IgG-mediated, but not direct DENV infection, these results suggest that TBC1D24 and SV2B have minimal roles in direct (non-ADE) DENV infection.

### 3.3. TBC1D24 and SV2B are required for efficient ADE in multiple contexts

We evaluated the role of TBC1D24 and SV2B in mediating ADE in various settings. First, to extend our validation studies with monoclonal antibodies above, we performed ADE assays in K562 cells using DENV2-GFP in the presence of serially diluted convalescent sera from three different DENV-immune donors (**Figure 3A**). TBC1D24 KO ablated ADE mediated by all three serum samples to similar levels seen with FcgRIIa KO control cells. SV2B KO also reduced ADE efficiency compared to WT cells, though its effect was more moderate compared to KO of TBC1D24 or FcgRIIa.

**Figure 3.**
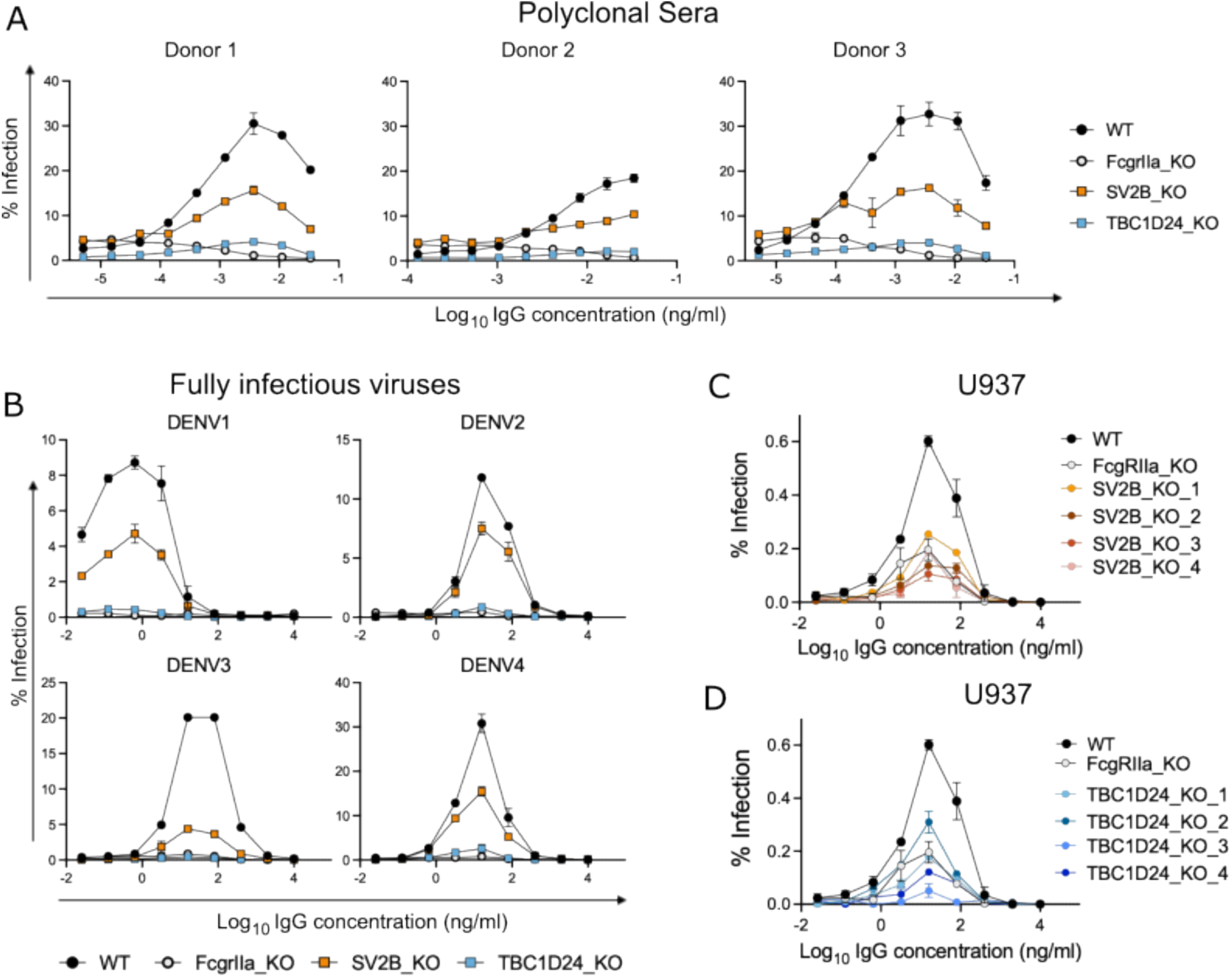
The role of TBC1D24 and SV2B in ADE is not limited to a single DENV serotype, antibody, or cell line. (**A**) DENV2-GFP was incubated with serial dilutions of convalescent serum from three independent DENV-immune donors prior to infection of the indicated K562 cells. Data points represent the mean and the error bars represent the range of infection in duplicate wells, respectively. Graphs shown are representative of two independent experiments. (**B**) Fully infectious DENV1-4 particles were incubated with serial dilutions of anti-DENV monoclonal IgG antibody (J9) prior to infection of the indicated K562 cells. Data points represent the mean and the error bars represent the range of infection in duplicate wells, respectively. Graphs shown are representative of two independent experiments. **(C-D)** DENV2-GFP was incubated with J9 monoclonal antibody prior to infection of clonal U937 cells with a KO in (**C)** SV2B or **(D)** TBC1D24. Data points represent the mean and the error bars represent the range of infection in duplicate wells. Data shown is representative of 4 independent experiments, each performed in duplicate wells. In each experiment, WT U937 cell pool and a U937 FcgRIIa KO clone were included as controls.

Next, to extend our findings with single-round DENV2-GFP, we performed ADE assays in K562 cells using fully infectious versions of all four DENV serotypes pre-complexed with J9 human monoclonal IgG (**Figure 3B**). ADE of all four DENV serotypes was abrogated in both TBC1D24-KO and FcgRIIa-KO cells. In contrast, SV2B KO reduced ADE of DENV1-4 to varying extents. Specifically, the strongest ADE reduction was observed for DENV3, and the weakest for DENV2 (∼80% and ∼30% reduction in peak infection, respectively). SV2B KO reduced peak enhancement of infection of both DENV1 and DENV4 by ∼50%.

Finally, we investigated the requirement for TBC1D24 and SV2B for efficient ADE in U937 cells (**Figure 3C-D**). Like K562 cells, U937 cells are commonly used to study ADE due to their susceptibility to efficient DENV infection only in the presence of IgG antibodies [^8,9^]. However, unlike K562 cells, U937 cells also express FcgRI in addition to FcgRIIa; both FcgRs are known to mediate ADE [^35^]. We tested four independent U937 KO clones each for TBC1D24 and SV2B (**Figure S8-9**) and included a previously generated U937 FcgRIIa KO clone [^35^] as a control in dose-response ADE assays. While KO of FcgRIIa abolished ADE in K562 cells (**Figures 3A-B**), it only partially reduced ADE efficiency in U937 cells (**Figures 3C-D**). The remaining ADE activity observed in FcgRIIa KO U937 is likely mediated by intact FcgRI expressed on these cells [^35^]. Notably, the reduction in ADE efficiency observed in SV2B-KO (**Figure 3C**) and TBC1D24-KO (**Figure 3D**) clones were comparable to, or in some cases (for TBC1D24 KO clones 3 and 4, **Figure 3D**), stronger than FcgRIIa KO in U937 cells.

Our combined results above show that the functional role of TBC1D24 and SV2B in ADE of DENV infection is not limited to a specific antibody, DENV serotype, or cell line.

### 3.4. TBC1D24 and SV2B facilitate binding of DENV-IgG complexes to cells

To determine at which step of the viral replication cycle TBC1D24 and SV2B are required during ADE, we first assessed the efficiency of binding and internalization of DENV2-IgG complexes into gene KO clones relative to WT K562 cells. Specifically, we measured cell-associated viral RNA levels by qRT-PCR either immediately following incubation of cells with IgG-virion complexes at 4°C (0h, binding), or following additional incubation at 37°C (internalization) for various time points. As expected, FcgRIIa KO cells substantially reduced cell-associated viral RNA levels at the initial timepoint, which corresponds to binding of IgG-DENV complexes to cells (79% reduction relative to WT, **Figure 4A**). Cell-associated viral RNA levels in TBC1D24-KO and SV2B-KO cells were also reduced at this initial timepoint (50% reduction for each compared to WT) (**Figure 4A**), indicating that TBC1D24 and SV2B promote binding of IgG-DENV complexes to cells.

**Figure 4.**
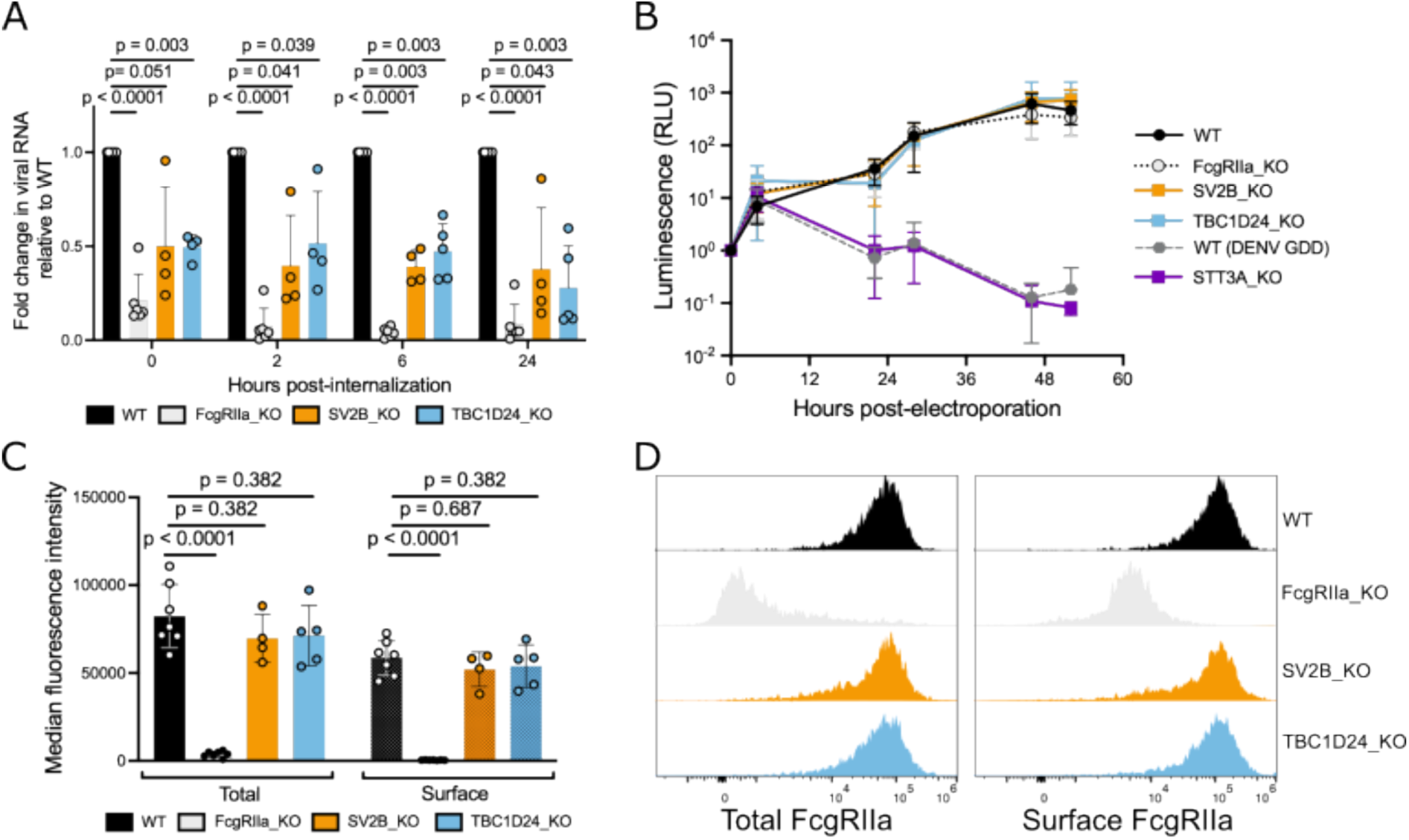
TBC1D24 and SV2B mediate efficient binding of DENV-IgG complexes to cells. (**A**) Quantitative RT-PCR of DENV RNA from cell-surface and internalized virions in K562 cells. DENV2-GFP pre-complexed with J9 monoclonal antibody were added to cells on ice for one hour. Cells were then washed to remove unbound virions and harvested either immediately (0h) or following additional incubation at 37C for the indicated time points (x-axis). Bars represent the mean normalized to WT at each time point from at least four independent experiments (data points) performed in duplicate wells and error bars show the standard deviation. **(B)** Relative luminescence of the indicated K562 cells electroporated with Renilla luciferase-expressing DENV2 replicon and lysed at indicated time points. Values for each cell line were normalized to the corresponding 0-hour time point to account for differences in electroporation efficiencies. Data points show the mean of four independent experiments, and error bars show the standard deviation. **(C)** Median fluorescence intensity of FcgRIIa expression in permeabilized (total expression) or unpermeabilized (surface expression) K562 cells. Bars represent the mean from at least four independent experiments (data points) and error bars show the standard deviation. **(D)** Histograms of FcgRIIa expression from a representative experiment of the data shown in (**C**). For (**A**) and (**C**), p-values shown are from multiple unpaired student’s t-tests adjusted using the Benjamini-Hochberg method.

Next, to confirm an impact on early infection steps, we performed an established luciferase replicon assay by electroporating DENV2-luciferase RNA into K562 cells to bypass entry [^38^]. Reporter gene activity over the first 10-12 h reflects translation of the input viral positive-stranded RNA genome while subsequent increases in signal are due to viral genome replication [^38^]. As a control, we included WT cells electroporated with replication-deficient mutant DENV (DENV GDD). We also included clonal K562 cells with a KO in STT3A, a host factor required for DENV RNA replication [^38^]. As expected, luminescence activity in replication-impaired controls (STT3A KO cells or WT cells electroporated with DENV GDD) were comparable to WT at early time points but were impaired beginning 23 hours post-electroporation (**Figure 4B**), indicating inhibition of viral genome replication, but not translation. Also as expected, FcgRIIa KO cells had no impact on viral genome translation or replication (**Figure 4B**). Reporter gene expression in TBC1D24 KO and SV2B KO cells mirrored that in WT and FcgRIIa KO cells across all time points, indicating limited effects on viral genome translation and replication (**Figure 4B**). Together, these results demonstrate that TBC1D24 and SV2B act on early stages of ADE, starting from binding of IgG-DENV complexes to cells.

Because we observed an impact on binding of IgG-DENV complexes to K562 cells, we asked whether KO of TBC1D24 or SV2B impaired expression of FcgRIIa, the sole FcgR expressed on these cells. Total and surface FcgRIIa expression was comparable between KO and unmutagenized WT K562 cells, as assessed by both median fluorescence intensity values (**Figure 4C**) and distribution of FcgRIIa expression across the cell population (**Figure 4D**). These results indicate that reduction in binding efficiency of DENV-IgG in TBC1D24-KO and SV2B-KO cells was not due to overt defects in FcgRIIa expression.

## 4. Discussion

By performing unbiased genome-wide and follow-up targeted CRISPR knockout screens, we identify for the first time candidate host dependency factors beyond FcgR that are exclusively required for ADE. Of these novel factors, we validated a functional role for TBC1D24 and SV2B in mediating efficient ADE of all four serotypes of DENV, using both monoclonal antibodies and polyclonal sera, and in multiple cell lines.

TBC1D24 and SV2B have established functions in trafficking specialized recycling endosomes relevant for neurotransmission (^43,44^). To our knowledge, a role for TBC1D24 in viral infection had not been described prior to our study. However, other TBC proteins, namely TBC1D16 and TBC1D20, have been shown to display antiviral and proviral activities, respectively [^78,79^]. Beyond its well defined role in synaptic vesicle trafficking, SV2 acts as the receptor for botulinum toxin [^80^]. Additionally, Sindbis virus infection upregulates expression of homologs of mammalian SV2 in *Aedes aegypti* mosquitoes [^81^], which are the primary vectors for DENV. Combined with our findings here, these studies suggest that viruses can subvert host factors involved in regulated secretion, unexpectedly including those that traditionally mediate neurotransmission. Indeed, Zika virus, a flavivirus closely related to DENV, enhances the expression of synaptotagmin-9 protein, a calcium sensor in neuroendocrine cells, and alters its subcellular localization [^82,83^]. Moreover, TMEM41B, which is involved in synaptic transmission in motor circuit neurons [^84,85^], is a host dependency factor for infection by multiple flaviviruses [^41^] and coronaviruses [^86–89^]. As described above, in the context of ADE of DENV infection, our targeted screens also identify other top hits with links to synaptic processes (**Figure 1E**), though their roles in ADE remain to be functionally validated. Nevertheless, their enrichment suggests that non-canonical IgG-mediated DENV entry [^21^] could exploit unconventional endocytic pathways described for synaptic processes [^90^].

Although the efficiency of ADE of DENV2-GFP mediated by monoclonal IgG was impaired to a similar extent by KO of either TBC1D24 or SV2B (**Figures 2A-B**), the former had a broader and more substantial impact in ADE assays using all four DENV serotypes and in the presence of polyclonal sera (**Figures 3A-B**). This finding implies a more critical role for TBC1D24 in ADE that is robust to assay conditions and may partly explain why SV2B was enriched in the genome-wide screen, but not the more stringent follow-up targeted screen. An alternative explanation is that in addition to its requirement for efficient ADE, SV2B may play a minor role in direct infection. Although the median reduction in direct infection efficiency of SV2B KO clones was comparable to control FcgRIIa KO clones, we note that two of four SV2B KO clones displayed a substantial reduction in direct infection efficiency compared to WT K562-DCSIGN cells. Further studies using additional clones will be required to clarify the relative contribution of SV2B to ADE and direct infection. As TBC1D24 and SV2B have shared functions in vesicle trafficking, it is also possible that they have partially redundant roles in ADE. This hypothesis can be tested in future studies examining whether ADE efficiency in SV2B KO cells can be rescued by overexpression of TBC1D24 and vice versa.

We found that KO of TBC1D24 and SV2B impaired efficient binding of IgG-bound DENV to K562 cells. Given the established role of FcgRIIa in mediating binding and internalization of IgG-DENV complexes [^14^], we were surprised that KO of TBC1D24 or SV2B minimally impacted FcgRIIa expression. As FcgRIIa association with lipid rafts has been shown to be important for ligand binding activity [^91–93^], it is possible that TBC1D24 and SV2B instead regulate the composition of cell membranes and/or trafficking of FcgRIIa to specific membrane microenvironments. Another possibility is that these host factors are involved in actin-driven cell membrane protrusions that have been implicated as a novel FcgR-dependent mechanism to capture IgG-bound DENV particles [^21^].

Another unexpected finding is the high enrichment of HELZ2 in both genome-wide and targeted screens. HELZ2 is an interferon-stimulated helicase and nuclear factor coactivator [^55,56,94^] with previously described *antiviral* activity in the context of direct DENV and to a lesser extent, Zika virus, infection [^55,56^]. Thus, HELZ2 may be among a growing set of interferon-stimulated genes with both antiviral and proviral functions [^55,95^]. HELZ2 appears to exert its anti-DENV activity by modulating host lipid metabolism following direct infection [^55^]. This finding raises a possible link between HELZ2 and one of the above hypothetical *proviral* mechanisms of TBC1D24 and SV2B in the context of ADE. There are two human isoforms of HELZ2, of which the longer isoform appears to exhibit higher interferon responsiveness [^55,56^]. Although both isoforms are targeted by the most enriched sgRNAs in our targeted screen, it remains to be determined whether the apparent proviral activity of HELZ2 in the context of ADE is isoform-dependent.

One limitation of our study is that although we validated the requirement of TBC1D24 and SV2B for efficient ADE in multiple cell lines, we were unable to confirm our findings in primary cells due to difficulty in maintaining cell viability following CRISPR editing and subsequent infection via ADE. Another limitation is that we were unable to detect TBC1D24 and SV2B in unedited WT cells via western blotting so we could not confirm loss of protein expression in KO cells. Notably, expression of cellular factors at levels insufficient for protein detection can nevertheless affect virus infection, as demonstrated for the alphavirus receptor, MXRA8 [^96^], and the interferon stimulated gene, LY6E [^97^]. Moreover, our ability to rescue ADE efficiency in KO cells via *trans*-complementation with the gene of interest supports a specific functional requirement for these host factors (**Figures 2A-B**). It is possible that infection with DENV via ADE upregulates the otherwise limited endogenous expression of TBC1D24 and SV2B proteins in non-neuronal cells.

In summary, our screen highlights features shared between direct infection and ADE, and those exclusively required during ADE. Among the latter, we demonstrated a functional role for TBC1D24 and SV2B in promoting efficient ADE of DENV infection in multiple contexts. TBC1D24 and SV2B were not enriched in previous genome-scale screens that reproducibly identified host dependency factors for direct flavivirus infection [^38–42^] and have no known roles in virus infection in general. In the absence of a biologically relevant *in vivo* model that can recapitulate dengue disease and immunity, this *in vitro* study is a key step in advancing our limited knowledge surrounding the biology of ADE of DENV. Further validation and mechanistic studies of TBC1D24, SV2B, and other screen hits can also lay the foundation for discovering host proteins and pathways that can be targeted by antiviral drugs to thwart dengue disease. Of note, SV2 proteins are the target of existing anti-epileptic drugs, some of which are FDA-approved [^98–101^]. It would be interesting to test the ability of these drugs to disrupt ADE processes.

## Author Contributions

Conceptualization, LG; Methodology, LB, MC, CHCA., LML, CDM, AA, JL, LG; Validation, LB, MC, CHC, LML, CDM, AA, JL, LG; Formal Analysis, LB, MC, CHCA, LML, CDM, AA, JL, LG; Investigation, LB, MC, CHCA, LML, CDM, AA, JL, LG.; Resources, LG; Writing – Original Draft Preparation, LB and LG; Writing – Review & Editing, LB, CHCA, CDM, LG. Visualization, LB, MC, CHCA, LML, JL, LG; Supervision, LG.; Project Administration, LG; Funding Acquisition, LG.

## Data Availability Statement

All pertinent data are within the manuscript and its supplement. Full sequencing data supporting this study are publicly accessible under GEO Accession number GSE264625.

## Acknowledgments

We thank Pritha Chanana (Fred Hutchinson Cancer Center Bioinformatics Shared Resources) for assistance with MAGeCK analysis and custom targeted library design; Fred Hutchinson Cancer Center Flow Cytometry (Rebecca Reeves, Nate Colven, Erik Huynh) and Antibody Technology (Norman Boiani, Catherina Artikis, Leticia Vasquez) Shared Resources; Ted Pierson for providing Raji-DCSIGNR cells; Andreas Puschnik for providing K562-Cas9-Blast cells; Michael Emerman for providing TZM-bl cells; Taia Wang for providing FcgRIIa KO U937 cells; Michael Diamond for providing DV2-70 antibody; Jan Carette for providing DENV2-luciferase replicon; Sandra Bajjalieh, Michael Emerman, Molly OHainle, and Jihong Bai for advice related to work on this manuscript; and Roberto Carlos Segura for assistance with data visualization and input on this manuscript. We obtained fully infectious DENV1-4 isolates and DENV-immune sera through BEI Resources, NIAID, NIH, as part of the World Reference Center for Emerging Viruses and Arboviruses program (WRCEVA).

