## Supplementary material for "Functional genomics screens reveal a role for TBC1D24 and SV2B in antibody-dependent enhancement of dengue virus infection": Figure S1

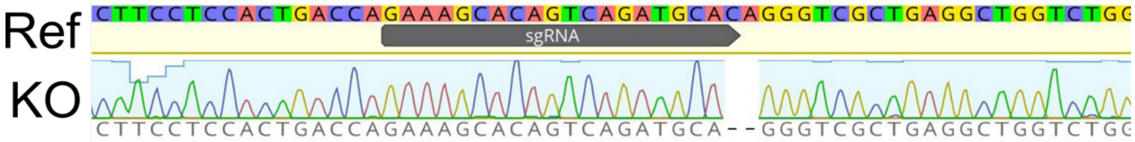


**Fig S1: Genotyping of K562 FcgRIIa KO clone.**
Sanger sequencing of locus targeted by sgRNA in the K562 FcgRIIa KO clonal line. Traces were aligned to WT reference sequence (“Ref”) to identify the indicated deletion.
