## Supplementary material for "Functional genomics screens reveal a role for TBC1D24 and SV2B in antibody-dependent enhancement of dengue virus infection": Figure S2

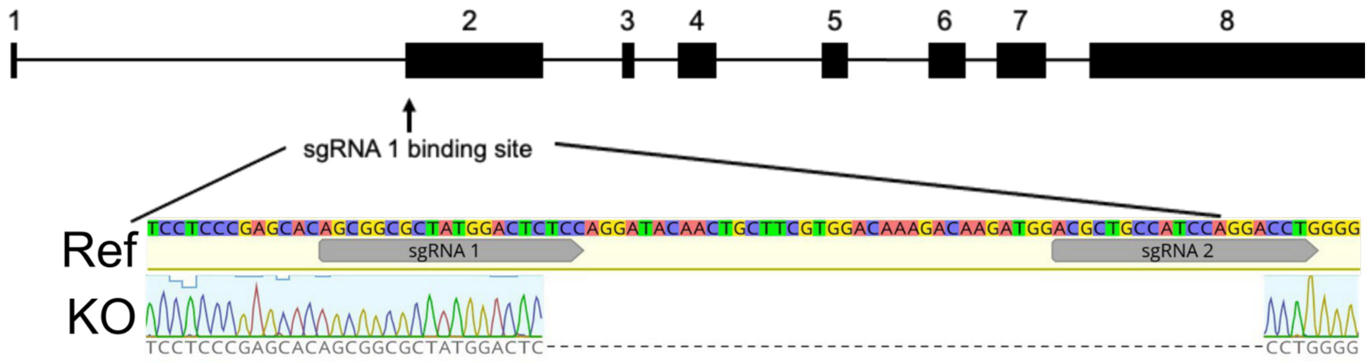


**Fig S2: Genotyping of K562 TBC1D24 KO clone.**
(Top) schematic of TBC1D24 exons (boxes) and introns (lines). (Bottom) Sanger sequencing of loci targeted by sgRNA1 (Table S4) in the K562 TBC1D24 KO clonal cell line. Traces were aligned to WT reference sequence (“Ref”) to identify the indicated deletion within exon 2.
