## Supplementary material for "Functional genomics screens reveal a role for TBC1D24 and SV2B in antibody-dependent enhancement of dengue virus infection": Figure S3

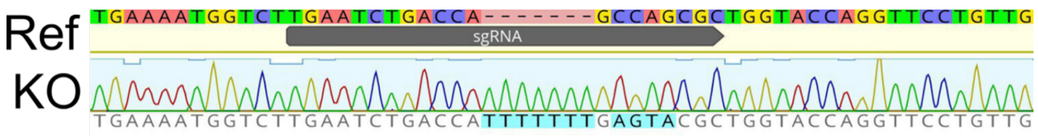


**Fig S3: Genotyping of K562 SV2B KO clone.**
Sanger sequencing of locus targeted by sgRNA in K562 SV2B KO clonal cell line. Traces were aligned to WT reference sequence (“Ref”) to identify a 7 bp insertion and 4 bp missense mutation.
