## Supplementary material for "Functional genomics screens reveal a role for TBC1D24 and SV2B in antibody-dependent enhancement of dengue virus infection": Figure S4

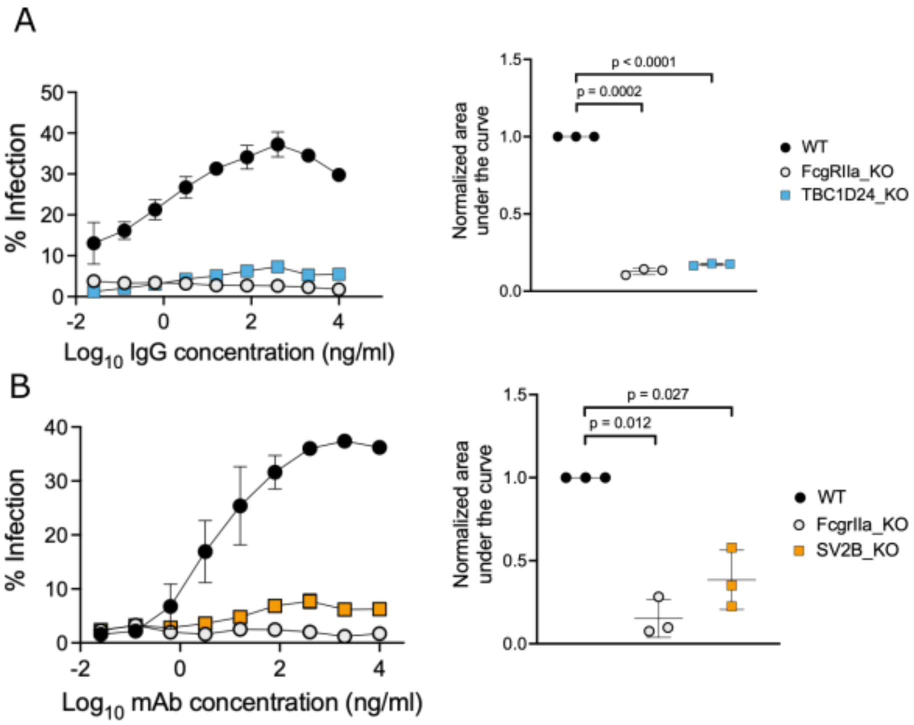


**Fig S4: Functional validation of TBC1D24 and SV2B in ADE assays**
**A-B**: (Left) The indicated K562 cells were infected via ADE using DENV2-GFP in the presence of serially diluted mouse anti-DENV IgG monoclonal antibody DV2-70 used in CRISPR screens. Data points represent the mean of three independent experiments normalized to the peak infection level of WT cells, and the error bars represent the standard deviation. (Right) Quantification of area under the curve normalized to WT K562 cells from three independent dose-response ADE experiments (data points), each performed in duplicate wells. Horizontal lines and error bars indicate mean and standard deviation, respectively. In each experiment, a FcgRIIa KO clone was included as a control. P-values shown are from multiple independent paired student's t-tests adjusted using the Benjamini-Hochberg method.
