## Supplementary material for "Functional genomics screens reveal a role for TBC1D24 and SV2B in antibody-dependent enhancement of dengue virus infection": Figure S5

| Clone | KO Score |
|---|---|
| KO_1 | 92 |
| KO_2 | nd |
| KO_3 | 97 |
| KO_4 | nd |

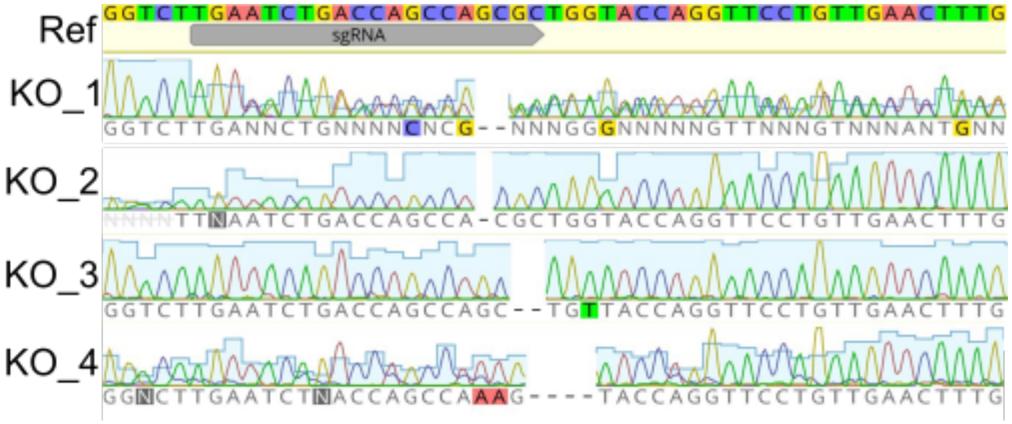


**Fig S5: Genotyping of K562-DCSIGN SV2B KO clones.**
(Right) Sanger sequencing of locus targeted by sgRNA in the K562-DCSIGN SV2B KO clones. Traces were aligned to WT reference sequence and heterogeneous mutations deconvoluted using Inference of CRISPR Edits (ICE; https://ice.synthego.com/#/). (Left) KO scores as determined by ICE; nd = not determined.
