## Supplementary material for "Functional genomics screens reveal a role for TBC1D24 and SV2B in antibody-dependent enhancement of dengue virus infection": Figure S7

| Clone | KO Score |
|---|---|
| KO_1 | 100 |
| KO_2 | 100 |
| KO_3 | 100 |
| KO_4 | 100 |
| KO_5 | 95 |
| KO_6 | 100 |
| KO_7 | 100 |
| KO_8 | 100 |
| KO_9 | 96 |
| KO_10 | 100 |
| KO_11 | 100 |

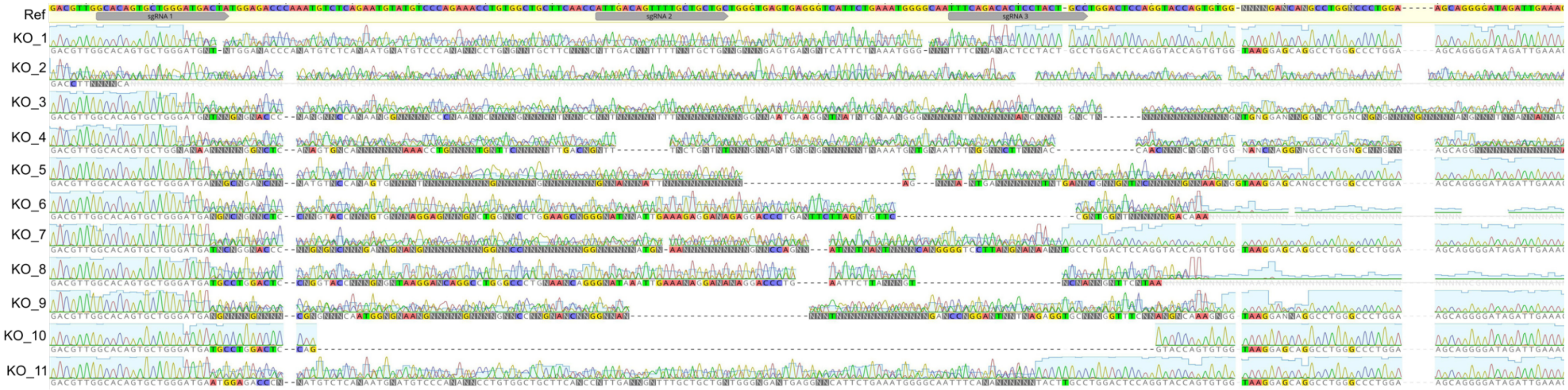


**Fig S7: Genotyping of K562-DCSIGN FcgRIIa KO clones**
(Right) Sanger sequencing of locus targeted by gRNA K562-DCSIGN FcgRIIa KO clones. Traces were aligned to WT reference sequence and heterogeneous mutations deconvoluted using ICE (https://ice.synthego.com/#/). (Left) KO scores as determined by ICE.
