## Supplementary material for "Functional genomics screens reveal a role for TBC1D24 and SV2B in antibody-dependent enhancement of dengue virus infection": Figure S9

| Clone | KO Score |
|---|---|
| KO_1 | 100 |
| KO_2 | 100 |
| KO_3 | 100 |
| KO_4 | 100 |

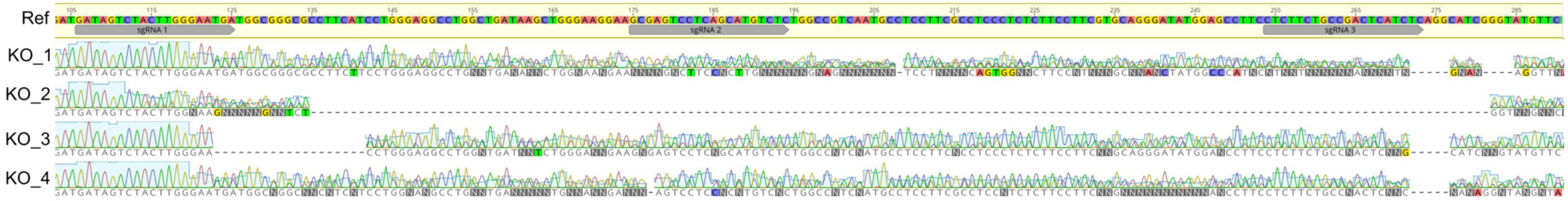


**Fig S9: Genotyping of U937 SV2B KO clones**
(Right) Sanger sequencing of locus targeted by gRNA in U937 SV2B KO clones. Traces were aligned to WT reference sequence and heterogeneous mutations deconvoluted using ICE (https://ice.synthego.com/#/). (Left) KO scores as determined by ICE.
